# Optimal Response Vigor and Choice Under Non-stationary Outcome Values

**DOI:** 10.1101/106500

**Authors:** Amir Dezfouli, Bernard W. Balleine, Richard Nock

## Abstract

Within a rational framework a decision-maker selects actions based on the reward-maximisation principle which stipulates they acquire outcomes with the highest values at the lowest cost. Action selection can be divided into two dimensions: selecting an action from several alternatives, and choosing its vigor, i.e., how fast the selected action should be executed. Both of these dimensions depend on the values of the outcomes, and these values are often affected as more outcomes are consumed, and so are the actions. Despite this, previous works have addressed the computational substrates of optimal actions only in the specific condition that the values of outcomes are constant, and it is still unknown what the optimal actions are when the values of outcomes are non-stationary. Here, based on an optimal control framework, we derive a computational model for optimal actions under non-stationary outcome values. The results imply that even when the values of outcomes are changing, the optimal response rate is constant rather than decreasing. This finding shows that, in contrast to previous theories, the commonly observed changes in the actions cannot be purely attributed to the changes in the outcome values. We then prove that this observation can be explained based on the uncertainty about temporal horizons; e.g., in the case of experimental protocols, the session duration. We further show that when multiple outcomes are available, the model explains probability matching as well as maximisation choice strategies. The model provides, therefore, a quantitative analysis of optimal actions and explicit predictions for future testing.

## Introduction

According to normative theories of decision-making, actions made by humans and animals are chosen with the aim of earning the maximum amount of future reward whilst incurring the lowest cost (Marshall, 1890; von Neumann & Morgenstern, 1947). Within such theories individuals optimize their actions by learning about their surrounding environment so as to satisfy their long-term objectives. The problem of finding the optimal action is, however, argued to have two aspects: (1) choice, i.e., deciding which action to select from several alternatives; and (2) vigor, i.e., deciding how fast the selected action should be executed. For a rat in a Skinner box, for example, the problem of finding the optimal action involves selecting a lever (choice) and deciding at what rate to respond on that lever (vigor). High response rates can have high costs (e.g., in terms of energy consumption), whereas a low response rate could have an opportunity cost if the experimental session ends before the animal has earned sufficient reward. Optimal actions provide the right balance between these two factors and, based on the reinforcement-learning framework and methods from optimal control theory, the characteristics of optimal actions and their consistency with various experimental studies have been previously elaborated (Dayan, 2012; Niv, Daw, Joel, & Dayan, 2007; Salimpour & Shadmehr, 2014).

These previous models have assumed, however, that outcome values are stationary and do not change on-line over the course of a decision-making session. To see the limitations of such an assumption, imagine the rat is in a Skinner box and has started to earn outcomes (e.g., food pellets) by taking actions. One can assume that, as a result of consuming rewards, the motivation of the animal to earn more food outcomes will decrease (e.g., because of satiety) and, therefore, over time, the outcomes earned will have a lower value. Such changes in value should affect both optimal choice and vigor (Killeen, 1995) but have been largely ignored in the previous models. This is while in most of the experimental and real-world scenarios, outcome values are affected by the history of outcome consumption, a phenomenon known as “law of diminishing marginal utility”*1* in the economics literature, and as “drive reduction theory” in psychological accounts of motivation, which indicates that the drive for earning an outcome decreases as the consequence of prior consumption of the outcomes (Hull, 1943; Keramati & Gutkin, 2014).

Here, building on previous work, we introduce a new concept called *reward field*, which captures non-stationary outcome values. Using this concept and methods from optimal control theory, we derive the optimal response vigor and choice strategy without assuming that outcome values are stationary. In particular, the results indicate that even when the values of outcomes are changing, the optimal response rate in an instrumental conditioning experiment is a constant response rate. This finding rules out previous suggestions that the commonly observed decrease in within-session response rates is due to decreases in outcome value (Killeen, 1995). Instead, we show that decreases in within-session response rates can be explained by uncertainty regarding session duration. This later analysis is possible because the session duration is explicitly represented in the current model, which is another dimension in which the current model extends previous work. The framework is then extended to choice situations and specific predictions are made concerning conditions under which the optimal strategy is maximization or probability matching.

## Model Specification

### The outcome space

We define the *outcome space* as a coordinate space with *n* dimensions, where *n* is the number of outcomes in the environment. For example, in a concurrent instrumental conditioning experiment in which the outcomes are water and food pellets, the outcome space will have two dimensions corresponding to water and food pellets. Within the outcome space, the state of the decision-maker at time *t* is defined by two factors: (i) the amount of *earned outcome* up to time *t*, which is denoted by **x**_*t*_and can be thought of as the position of the decision-maker in outcome space; e.g., in the above example, **x**_*t*_ = [1,2] would indicate that one unit of water and two units of food pellet have been gained up to time *t*; and (ii) the *outcome rate* at time *t*, denoted by **v**_*t*_, which can be considered the velocity of the decision-maker in the outcome space 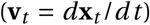; e.g., if a rat is earning two units of water and one unit of food pellet per unit of time, then **v**_*t*_ = [2,1]. In general, we assume that the outcome rate cannot be negative (**v** ≥ 0), which means that the cumulative number of earned outcomes cannot decrease with time.

### The reward

We assume that there exists an *n*-dimensional *reward field*, denoted by *A*_**x**, *t*_, where each element of *A*_**x**, *t*_ represents the per unit value of each of the outcomes. For example, the element of *A*_**x**, *t*_ corresponding to food pellets represents the value of one unit of food pellet at time *t*, given that **x** units of outcome have been previously consumed. As such, *A*_**x**, *t*_ is a function of both time and the amount of outcome earned. This represents the fact that (i) the reward value of an outcome can change value as a result of consuming previous outcomes, e.g., due to satiety (depending on **x**) and (ii) the reward value of an outcome can change purely with the passage of time; e.g., an animal can get hungrier over time causing the reward value of food pellets to increase (depending on *t*). In general, we assume that *A*_**x**, *t*_ has two properties:

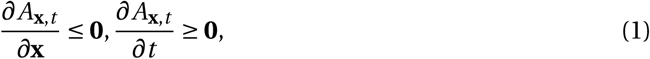
which entail that (i) the outcome values decrease (or remain constant) as more outcomes are earned, and (2) that outcome values do not decrease with the passage of time.

### Cost

Within the context of instrumental conditioning experiments, previous studies have expressed the cost of earning outcomes as a function of the delay between consecutive responses made to earn outcomes. For example, if a rat is required to make several lever presses to earn outcomes, then the cost will be higher if the delay between lever presses is short. More precisely, if the previous response has occurred *τ* time steps ago, then the cost of the current lever press will be (Dayan, 2012; Niv et al., 2007):

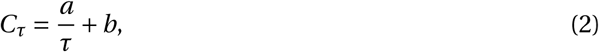
where *a* and *b* are constants. *b* is the constant cost of each lever press, which is independent of the delay between lever presses whereas the factor *a* controls the rate-dependent componentof the cost. Previous research has established that predictions derived from this definition of cost are consistent with experimental data (Dayan, 2012; Niv et al., 2007). Note that costs such as basal metabolic rate and the cost of operating the brain, although consuming a high portion of energy produced by the body, are not included in the above definition because they are constant and independent of response rate and, therefore, are not directly related to the analysis of response vigor and choice.

Here, we express cost as a function of outcome rate rather than the rate of action execution. We define the cost function *K*_**v**_ as the cost paid at each time step for earning outcomes at rate **v**. In the specific case that the outcome space has one dimension (there is only one outcome), and under ratio schedules of reinforcement (fixed-ratio, variable-ratio, random-ratio) in which the decision-maker is required to perform either precisely or on average *k* responses to earn one unit of outcome, the cost defined in equation 2 will be equivalent to:

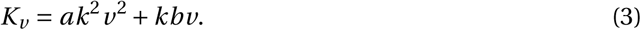

See Theorem A1 in Appendix for the proof. This definition of cost implies that it is only a function of outcome rate and is time-independent 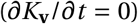. Although, in general, it may seem reasonable to assume that, as time passes within a session, the cost of taking actions will increase because of factors such as effector fatigue, here we made a time-independence assumption based on previous studies showing that factors such as effector fatigue have a negligible effect on response rate (McSweeney, Hinson, & Cannon, 1996). In general, we assume that at least one response is required to earn an outcome (*k* > 0), and the cost of earning outcomes is non-zero (*a* > 0).

### Value

The reward earned in each time step can be calculated as the reward of one unit of each of the outcomes (*A*_**x**, *t*_) multiplied by the amount earned from each of the outcomes, which will be **v**.*A*_**x**, *t*_. The cost of earning this amount of reward is *K*_**v**_, and therefore the net amount of reward earned will be:

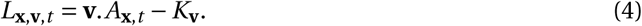

A decision-making session starts at *t* = 0 and the total duration of that session is *T*. We denote the total reward gained in this period as *S*_0, *T*_, which is the sum of the net rewards earned at each point in time:

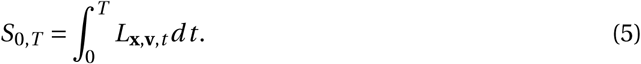

The quantity *S*_0, *T*_ is called the *value* function, and the goal of the decision-maker is to find the optimal rate of earning outcomes that yields the highest value. The optimal rates that maximize *S*_0, *T*_can be found using different variational calculus methods, such as the Euler-Lagrange equation or the Hamilton-Jacobi-Bellman equation (Liberzon, 2011). The results presented in the next sections are derived using the Euler-Lagrange equation (see Appendix for details of value function in non-deterministic schedules).

## Results

### Optimal response vigor

In this section we use the model presented above to analyse optimal response vigor when there is one outcome and one response available in the environment. The analysis is divided into two sections. In the first section, we assume that the decision-maker is certain about session duration, i.e., that the session will continue for *T* time units, and will extend this analysis in the next section to a condition in which the decision-maker assumes a probabilistic distribution of session lengths.

**Response vigor when the session duration is known.** We maintain the following theorem:

#### Theorem 1

If the duration of the session is fixed and the time-dependent change in the reward field is zero 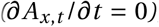, then the optimal outcome rate is constant 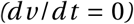. If the time-dependent change in the reward field is positive 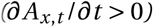, then the optimal outcome rate will be accelerating 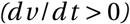.

See Appendix for a proof of this theorem. Note that the assumption 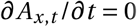 implies that the passage of time has no significant effect on the reward value of the outcome; e.g., a rat is not getting hungrier during an instrumental conditioning session, which is a reasonable assumption given the short duration of such experiments (typically less than an hour). Within this condition, the above theorem states that the optimal response rate is constant throughout the session, even under conditions in which the reward value of the outcome decreases within the session as a result of earning outcomes, e.g., because of satiety. As an intuitive explanation for why a constant rate is optimal, imagine a decision-maker who chooses a non-constant outcome rate that results in a total of *x*_*T*_ outcomes during the session. If, instead of the non-constant rate, the decision-maker selects a constant rate 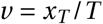, then the total outcomes earned will be the same as before; however, the cost will be lower because cost is a quadratic function of the outcome rate and, therefore, it is better to earn outcomes at a constant rate. Nevertheless, although this prediction is clear enough, it is not consistent with the experimental results, described next.

**
*Within-session pattern of responses.***
 It has been established that in various schedules of reinforcement, including variable-ratio (VR) schedules (McSweeney, Roll, & Weatherly, 1994), the rate of responding within a session has a particular pattern: the response rate reaches its maximum a short time after the session starts (warm-up period), and then gradually decreases toward the end of the session (Figure 1: left panel). Killeen (Killeen, 1994) proposed a mathematical description of this phenomenon, which can be expressed as follows (Killeen & Sitomer, 2003):

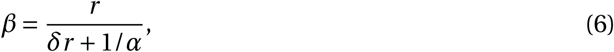
where *β* is the response rate, *δ* is the minimum delay between responses, *r* is the resulting outcome rate, and *α* is called *specific activation*^2^. The model suggests that as the decision-maker earns outcomes during the session, the value of *α* gradually declines due to satiety, which will Cause a decrease in response rate^3^. Although this model has been shown to provide a quantitative match to the experimental data, it is not consistent with Theorem 1 which posits that, even under conditions in which outcome values are changing within a session, the optimal response rate should not decrease during the session. As a consequence, the present model suggests that the cause of any decrease in the within-session response rate cannot be due purely to a change in outcome value.

Note, however, the optimal response rate advocated by Theorem 1 is not consistent with reports of decreasing response rates across a session, which implies that some of the assumptions made to develop the model may not be accurate. Although the form of the cost and reward functions is reasonably general, we assumed that the duration of the session, *T*, is fixed and known by the decision-maker. In the next section we show that relaxing this assumption such that the duration of the session is unknown results much closer concordance between predictions from the model and the experimental data.

**Figure 1.**
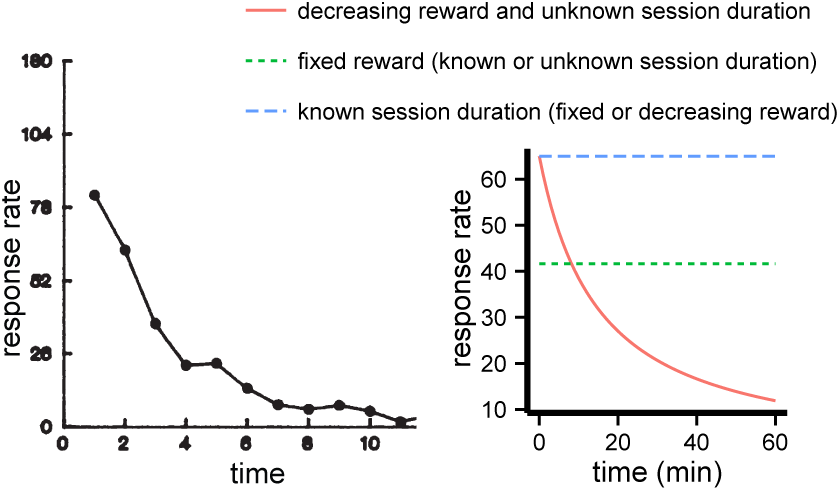
The pattern of within-session response rates (responses per minute). **Left panel**: Experimental data. The rate of responding per minute during successive intervals (each interval is 5 minutes) in a variable-ratio (VR15) schedule (*k* = 15). The figure is adopted from (McSweeney et al., 1994). **Right panel**: The theoretical pattern of within-session responses predicted by the model in different conditions. Please see the text for details of each condition.

**Response vigor when session duration is unknown. In** this section we assume that the decision-maker is uncertain about the session duration, which can be either because the session duration is in fact non-deterministic, or because of inherent inaccuracies in interval timing in animals (Gallistel & Gibbon, 2000; Gibbon, 1977). In this condition, a plausible way to calculate optimal response rates is to set an expectation as to how long the session will last and to calculate the optimal response rate based on that expectation. Based on this, if t^
*′*^ time step has passed since the beginning of the session, then the expected session duration is 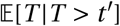 and therefore the value of the rest of the session will be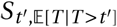. The following theorem maintains that the optimal rate of outcome delivery that maximizes the value function is a decreasing function of the current time in the session t^
*′*^, and therefore the optimal response rates will decrease throughout the session.

#### Theorem 2

Assuming 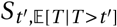 is the value function and that (i) the time dependent change in the reward field is zero 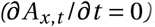, (ii) the probability that the session ends at each point in time

^3^Heresatiety refers to both post-ingestive factors (such as blood glucose level (Killeen, 1995)) and/or pre-ingestive factors (for example_-_sensory specific satiety: (McSweeney, 2004)).

Is non-zero (p(T) > 0), (iii) values of outcomes decrease as more are consumed 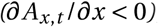, then the optimal rate of outcome delivery is a decreasing function of t^′^:

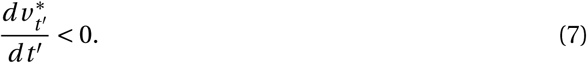

See Appendix for the proof of this theorem. Theorem 2 stems from two bases: (i) the optimal rate of outcome delivery is a decreasing function of session length, i.e., when the session duration is long the decision-maker can afford to earn outcomes more slowly, and (ii) when the session duration is unknown, expected session duration should increase with the passage of time. This phenomenon, which has been elaborated within the context of delayed gratification (McGuire & Kable, 2013; Rachlin, 2000), is more significant if the decision-maker assumes a heavy-tail distribution over *T*. Putting (i) and (ii) together implies that the optimal response rate will decrease with the passage of time. Importantly, this suggests, from a normative perspective, uncertainty about the session duration is necessary in order to explain within-session decreases in response rates.

For simulation of the model we characterized the session duration using a Generalized Pareto distribution following (McGuire & Kable, 2013). Simulations of the model are depicted in Figure 1: right panel. Simulations show three different conditions. In condition (i) the session duration is known, and as the figure shows irrespective of whether the reward of outcomes is decreasing or fixed, the optimal response rate is constant. In condition (ii) session duration is unknown, but the reward of outcomes is constant. Again in this condition the optimal response rate is constant. In condition (iii) session duration is unknown *and* the reward decreases as more outcomes are consumed. Only in this condition, consistent with experimental data, response rates decrease as time passes (see Appendix for details of the simulations). Therefore, the simulations confirm that decreases in the reward of outcomes alone are not sufficient to explain within-session response rates, but by assuming uncertainty about session duration, the pattern of responses will be consistent with the experimental data. Note that a similar pattern will be obtained using any other distribution that assigns a non-zero probability to positive values of *T*.

**
*Relationship to temporal discounting.***
There are, however, alternative explanations available based on changes in outcome value. One candidate explanation is based on the temporal discounting of outcome value according to which the value of future rewards is discounted compared to more immediate rewards. Typically, the discount value due to delay is assumed to be a function of the interaction of delay and outcome value. If, at the beginning of the session, outcome values are large (e.g., because a rat is more hungry), then any discount produced by selecting a slow response rate will be larger at that point than later in the session when the value of the outcome is reduced (e.g., due to satiety) and so a delay will have less impact. It could be argued, therefore, that it is less punitive to maintain a high response rate at the beginning of the session to avoid delaying outcomes and then to decrease response rate as time passes within the session. As such, temporal discounting predicts decreases in within-session response rates consistent both with experimental observations and with the argument that outcome value decreases within the session (e.g., the satiety effect).

**
*Prediction.***
Although plausible, such explanations make very different predictions compared to the model. The most direct prediction from the model is that introducing uncertainty into the session duration without altering the average duration should nevertheless lead to a sharper decline in response rate within the session; e.g., if for one group of subjects the session lasts exactly 30 minutes whereas for another group the session length is uncertain and can end at any time (but ends on average after 30 minutes), then the model predicts that the response rate in the second group will be higher at the start and decrease more sharply than in the first group. This effect is not anticipated by the temporal discounting account of the effect.

***Effect of experimental parameters: Relating the model to currently available data.*** Optimal response rates predicted by the model are affected by experimental parameters (e.g., reward magnitude), which can be compared against experimental data. In general, in an instrumental conditioning experiment, the duration of the session can be divided into three sections: (i) outcome handling/consumption time, which refers to the time that an animal spends consuming the outcome, (ii) *post-reinforcer pause*, which refers to the pause that occurs after consuming the outcome and before starting to make the next response (e.g., lever press). Such a pause is consistently reported in previous studies using an FR schedule, (iii) *run time*, which refers to the time spent making responses (e.g, lever pressing). Experimental manipulations have been shown to have different effects on the duration of these three sections of the session, and whether each of these sections is included when calculating response rates can affect the results. The predictions of the current model are with regard to response rate; whether manipulating experimental parameters are expected to change the two other measures (outcome handling and post-reinforcer pause) cannot be predicted by the current model. In the following sections, we briefly review the currently available data from instrumental conditioning experiments and their relationship to predictions of the model^4^.

**
*The effect of response cost (***
*a*
**
*and***
*b*

).
Experimental studies in rats working on a FR schedule (Alling & Poling, 1995) indicate that increasing the force required to make responses causes increases in both inter-response time and the post-reinforcement pause. The same trend has been reported in Squirrel monkeys (Adair & Wright, 1976). Consistent with this observation, the present model predicts that increases in the cost of responding, for example by increasing the effort required to press the lever (increases in *a* and *b*), lead to a lower rate of earned outcomes and a lower rate of responding (Figure 2). The reason for this is that, by increasing the cost, the response rate for each outcome should slow in order to compensate for the increase in the cost and so maintain a reasonable gap between the reward and the cost of each outcome.

**Figure 2.**
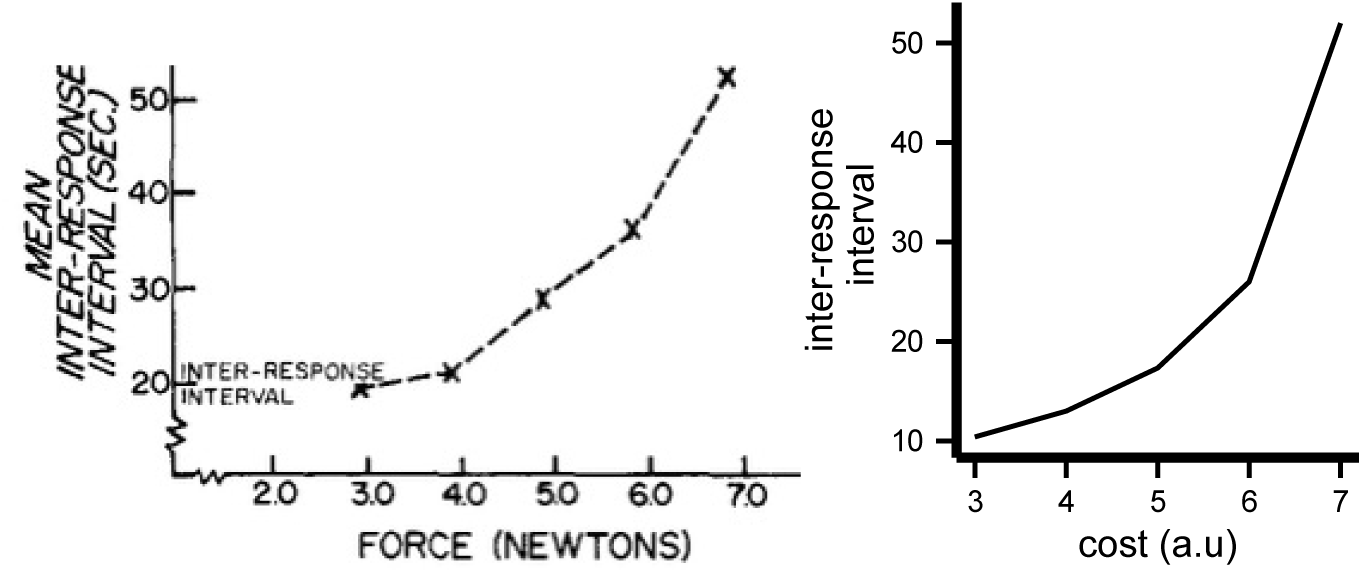
Effect of response cost on response rates.**Left panel**: Empirical data. Inter-response intervals when the force required to make a response is manipulated. Figure is adopted from Adair and Wright (1976). **Right panel**: Model prediction. Inter-response interval (equal to the inverse of response rates) as a function of cost of responses (*b*).

***The effect of ratio-requirement (****k*). Experimental studies mainly imply that the rate of earned outcomes decreases with increases in the ratio-requirement (Aberman & Salamone, 1999; Barofsky & Hurwitz, 1968), which is consistent with the general trend of the optimal rate of outcome earning implied by the present model (see below).

Experimental studies on the rate of responding on FR schedules indicate that the post-reinforcer pause increases with increases in the ratio-requirement (Ferster & Skinner, 1957, Figure 23)(Felton & Lyon, 1966; Powell, 1968;(Premack, Schaeffer, & Hundt, 1964) In terms of overall response rates, some experiments report that response rates increase with increases in the ratio-requirement up to a point beyond which response rates will start to decline, in rats (Barofsky & Hurwitz, 1968; Kelsey & Allison, 1976; Mazur, 1982), pigeons (Baum, 1993) and mice (Greenwood, Quartermain, Johnson, Cruce, & Hirsch, 1974), although other studies have reported inconsistent results in pigeons (Powell, 1968), or a decreasing trend in response rate with increases in the ratio-requirement (Felton & Lyon, 1966;(Foster, Blackman, & Temple, 1997). The inconsistency is partly due to the way that response rates are calculated in the different studies; for example 4Note that, for simplicity, the simulations in this section are made under the assumption that the session duration is fixed. In some studies outcome handling and consumption time are not excluded when calculating response rates (Barofsky & Hurwitz, 1968), in contrast to the other studies (Foster et al., 1997). As a consequence, the experimental data regarding the relationship between response rate and the ratio-requirement is inconclusive.

With regard to this issue, the present model predicts that the relationship between response rate and the ratio-requirement is an inverted U-shaped pattern (Figure 3: left panel), which is consistent with the studies mentioned previously depending on the value of other experimental parameters. The model makes an inverted U-shaped prediction because, under a low ratio-requirement, the cost is generally low and, therefore, as the ratio-requirement increases, the decision-maker will make more responses to compensate for the drop in the amount of reward. In contrast, when the ratio-requirement is high, the cost of earning outcomes is high and the margin between the cost and the reward of each outcome becomes significantly tighter as the ratio-requirement increases. The margin can, however, be loosened by decreasing the response rate.

**Figure 3.**
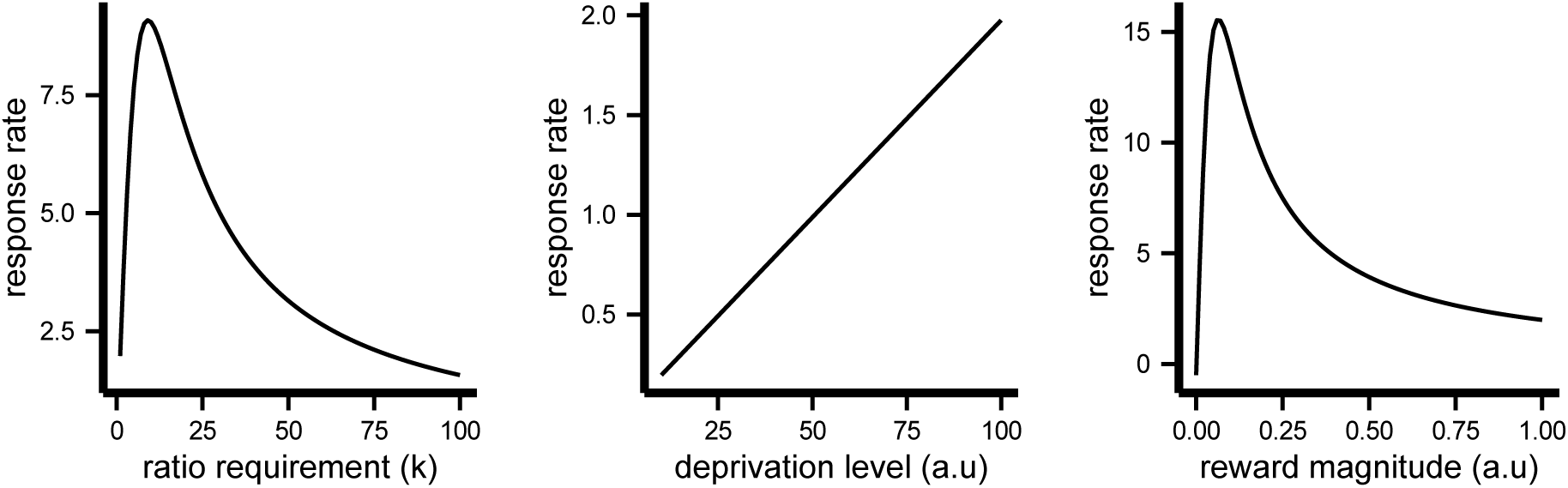
Left panel:The effect of ratio-requirement on the response rate. **Middle panel**: The effect of the initial motivational drive on response rates. **Right panel**: The effect of the reward magnitude on response rates.

**
*The Effect of deprivation level.***
Experimental studies that have used FR schedules suggest that response rates increase with increases in deprivation (Ferster & Skinner, 1957, Chapter 4)(Sidman & Stebbins, 1954). However, such increases are mainly due to decreases in the post-reinforcement pause, and not due to the increases in the actual rate of responding after the pause (see (Pear, 2001, Page 289) for a review of other reinforcer schedules). The model predicts that, with increases in deprivation, the rate of responding and of earned outcomes will increase linearly (Figure 3: middle panel). The rate of increase is, however, negligible when the session duration is long, in which case, even under high deprivation, sufficient time is available to earn sufficient reward and become satiated. This provides a potential reason why the effect of deprivation on response rate has not previously been observed in experimental data.

**
*The effect of reward magnitude.***
Some studies show that post-reinforcement pauses increase as the magnitude of the reward increases (Powell, 1969), whereas other studies suggest that the post-reinforcement pause decreases (Lowe, Davey, & Harzem, 1974), although, in this later study, the magnitude of the reward was manipulated within-session and a follow-up study showed that, at a steady state, the post-reinforcement pause increases with increases in the magnitude of the reward (Meunier & Starratt, 1979). Reward magnitude does not, however, have a reliable effect on the overall response rate (Keesey & Kling, 1961; Lowe et al., 1974; Powell, 1969). Regarding predictions from the model, the effect of the reward magnitude on earned outcome and response rates is, again, predicted go take an inverted U-shaped relationship (Figure 3: right panel), and, therefore, depending on the value of the parameters, the predictions of the model are consistent with the experimental data. The model makes a U-shaped prediction because, when the reward magnitude is large then, given high response rates, the animals will become satiated quickly and, therefore, the reward value of future outcomes will decrease substantially if animal maintains this high response rate. As a consequence, under a high reward magnitude condition, increase in reward will cause response rates to decrease. Under a low reward magnitude condition, however, satiety has a negligible effect and a high response rate ensures that sufficient reward can be earned before the session ends.

### Optimal choice and response vigor

In this section we address the choice problem, i.e., the case where there are multiple outcomes available in the environment and the decision-maker needs to make a decision about the response rate along each outcome dimension. An example of this situation is a concurrent instrumental conditioning experiment in which two levers are available and pressing each lever produces an outcome on a ratio schedule. Unlike the case with single outcome environments, the optimal rate of earning outcomes is not necessarily constant and can take different forms depending on whether the reward field is a *conservative field* or a *non-conservative field*, and whether the costs of responses along the outcome dimensions are independent of each other. Below, we derive the optimal choice strategy in each condition.

**
Conservative reward field.** A reward field is conservative if the amount of reward experienced by consuming different outcomes does not depend on the *order* of consumption and depends only on the number of each outcome earned by the end of the session. This property holds in two conditions (i) when the value of each outcome is unrelated to the prior consumption of other outcomes; and (ii) the consumption of an outcome affects the value of other outcomes and this effect is symmetrical. As an example of condition (i), imagine an environment with two outcomes in which one of the outcomes only satisfies thirst and the other only satisfies hunger. Here, consumption of one of the outcomes will not affect the amount of reward that will be experienced by consuming the other outcome and, therefore, the total reward during the session does not depend on the order of choosing the outcomes. As an example of condition (ii), imagine an environment with two outcomes in which both outcomes satisfy hunger and, therefore, consuming one of the outcomes reduces the amount of future reward produced by the other outcome. Here, if the effect of the outcomes on each other is symmetrical, i.e., consuming outcome O_1_, reduces the reward elicited by outcome O_2_ by the same amount that consuming outcome O_2_ reduces the reward elicited by outcome O_1_, then it will not matter which outcome is consumed first and the total reward during the session will be independent of the chosen order. As such, the reward field will be conservative.

More precisely, a reward field is conservative if there exists a scalar field *D*_**x**_ such that:

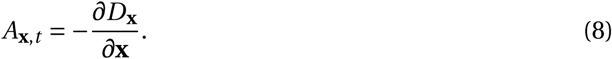

It can be shown that if a reward field satisfies equation 8 then the amount of reward experienced in a session depends on the total number of earned outcomes. Under this condition the optimal response rate will be constant for each outcome relative to the other. Intuitively, this is because, in terms of the total rewards per session, the only thing that matters is the final number of earned outcomes and, therefore, there is no reason why the relative allocation of responses to outcomes should vary within the session. The actual response rate for each outcome will, however, depend on whether the costs of the outcomes are independent, a point elaborated in the next section.

**
*Conservative reward field and independent response cost.***
The costs of various outcomes are independent if the decision-maker can increase their work for one outcome without affecting the cost of other outcomes. As an example, imagine a decision-maker that is using their left hand to make responses that earn one outcome and their right-hand to make responses that earn the second outcome. In this case, the independence assumption entails that the cost of responding with one or other hand is determined by the response rate on that hand; e.g., the decision-maker can increase or decrease rate of responding on the left hand without affecting the cost of responses on the right hand. More precisely, the independence assumption entails that the Hessian matrix of *K*_**v**_ is diagonal:

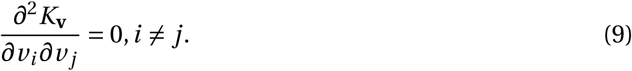

In this situation even if some of the outcomes have a lower reward or a higher cost (inferior outcomes) compared to other outcomes (superior outcomes), it is still optimal to allocate a portion of responses to the inferior outcomes. This is because responding for inferior outcomes has no effect on the net reward earned from superior outcomes and, therefore, as long as the response rate for inferior outcomes is sufficiently low that the reward earned from them is more than the cost paid, responding for them is justified. The portion of responses allocated to each outcome depends, however, on the cost and reward of each outcome. We maintain the following theorem:

#### Theorem 3

If (i) the reward field is conservative, i.e., there exists a scalar field D_**x**_ such that equation 8 is satisfied, (ii) the time-dependent term of the reward field is zero 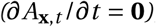, and (iii) the cost function satisfies equation 9, then the optimal rate of earning outcome **v\*** will be constant 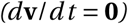 and satisfies the following equation:

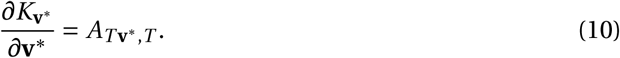

See Appendix for the proof and for the specification of optimal responses. As an example, imagine a concurrent fixed-ratio (FR) schedule in which an animal needs to make *k* responses on the left lever in order to earn O_1_, and *l k* responses on the right lever in order to earn O_2_, and both outcomes have the same reward properties. According to Theorem 3, the optimal response rate for O_1_ (the outcome with the lower ratio-requirement) is *l* times greater than the response rate for the second outcome O_2_. Figure 4: left panel independent cost condition shows the simulation of the model and the optimal trajectory in the outcome space. As the figure shows, the rate of earning O_1_ is higher than O_2_, however, the relative portion of earned outcomes remains the same throughout the session.

**Figure 4.**
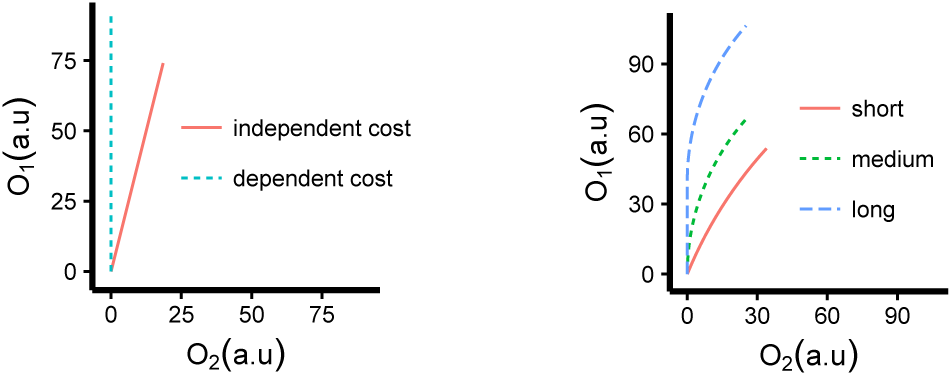
**Left panel**: Optimal trajectory in a conservative reward field. Earning **O**_1_ requires *k* responses and earning **O**_2_ requires *lk* responses. Initially the amount of earned outcome is zero (starting point is at point [0,0]), and the graph shows the trajectories that the decision-maker takes in two different conditions corresponding to when the costs of outcomes are independent, and when the costs are dependent on each other. **Right panel**: The optimal trajectories in the outcome space when the reward field is non-conservative. The graph shows the optimal trajectory in the conditions that the session duration is short (*T* = 7), medium (*T* = 15.75) and long (*T* = 23).

The above results are generally in line with the *probability matching* notion, which states that a decision-maker allocates responses to outcomes based on the ratio of responses required for each outcome (Estes, 1950). Probability matching is often argued to violate rational choice theory because, within this theory, it is expected that a decision-maker works exclusively for the outcome with the higher probability (i.e., the lower ratio-requirement). However, based on the model proposed here probability matching is the optimal strategy and therefore consistent with rational decision-making.

**
*Conservative reward field and dependent response cost.***
In this section we assume that the cost of responses for an outcome is affected by the rate of responses for earning other outcomes. In other words, what determines the cost is the delay between subsequent responses either for the same or for a different outcome; i.e., the cost is proportional to the rate of earning all of the outcomes. In instrumental conditioning this assumption entails, for example, that the cost of pressing, say, the right lever is determined by the time that has passed since the last press on either the right or a left lever. In this condition the optimal strategy is *maximisation*; i.e., to take the action with the higher reward (or lower ratio-requirement) and to stop taking the other action (see Theorem A2 in Appendix). The reason is, unlike the case with independent costs, allocating more responses to earn an inferior outcome will increase the cost of earning superior outcomes and, therefore, it is better to pay the cost for the superior outcome only, which requires fewer responses per unit of outcome.

For example, under a concurrent FR schedule in which an animal needs to make *k* responses on the left lever to earn O_1_, and *l k* responses on the right lever to earn O_2_ (O_1_ and O_2_ have the same reward properties), the optimal response rate will be a constant response rate on the left lever and a zero response rate on the right lever. Figure 4: left panel dependent cost condition shows a simulation of the model and the optimal trajectory in outcome space, which shows that the subject only earns O_1_.

As such, whether the outcome rate reflects a probability matching or a maximization strategy depends on the cost function and, in instrumental conditioning experiments, the cost that reflects the maximization strategy is more readily applicable. Regarding the experimental data, evidence from concurrent instrumental conditioning experiments in pigeons tested using VR schedules (Herrnstein & Loveland, 1975) is in-line with the maximization strategy and consistent with predictions from the model. Within the wider scope of decision-making tasks, some studies are consistent with probability matching, whereas other studies provide evidence in-line with the maximization strategy (see (Vulkan, 2000) for a review). In many of these latter studies, however, the decision-making task involved making a single choice (e.g., single button press) with immediate feedback (about whether the choice was rewarded) after which the next trial was initiated and, because such disjointed actions are unlikely to convey a rate-dependent cost, the structure of such studies cannot be readily related to the model proposed here.

**
*Prediction.***
One way of testing the above explanation for maximization and matching strategies is to compare the pattern of responses when two different effectors are used to make responses for the outcomes vs. when a single effector is being used to earn both outcomes. In the first condition the costs of the two outcomes are independent of each other whereas in the second condition they are dependent on each other. As a consequence, the theory predicts that, in the first condition, response rates will reflect probability matching whereas in the second condition they will reflect the maximization strategy.

**Non-conservative reward field.** A reward filed is non-conservative if the total amount of reward experienced during the session depends on the order of the consumption of the outcomes. Imagine an environment with two outcomes say O_1_ and O_2_, where both outcomes have the same motivational properties, e.g., consumption of one unit of either O_1_ or O_2_ decreases hunger by one unit, however, they generate different amount of rewards, e.g., one unit of O_1_ generates more reward than one unit of O_2_. Within such an environment, if hunger is high then consuming O_1_ generates significantly more reward than O_2_ and, therefore, early in the session it is better to allocate more responses to O_1_, whereas later in the session (when hunger is presumably lower and the difference in the value of the outcomes is small) responses for O_2_ can gradually increase. If we reverse this order, i.e., first O_2_ is consumed and then O_1_, then early consumption of O_2_ will cause satiety and the decision-maker will lose the chance to experience high reward from O_1_ when hungry. As such, the amount of experienced reward depends on the order of consuming the outcomes and, based on the above explanation, a larger amount of reward can be earned during the session if more responses are allocated to the outcome with the higher reward at the beginning of the session (see Theorem A3 in Appendix). Figure 4: right panel shows the simulations of the model under different session durations. In each simulation, at the beginning of the session the initially earned outcomes are zero and each line in the figure shows the trajectory of the amount earned from each outcome during the session. As the figure shows, in all conditions the rate of earning O_1_ is higher than O_2_ and this difference is more apparent under long session durations, in which a large amount of reward can be gained during the session and it makes a significant difference to earn O_1_ first.

**
*Prediction.***
 A test of the above prediction would involve an experiment in which the subject is responding for two food outcomes containing an equal number of calories (and therefore having the same impact on motivation) but with different levels of the desirability (e.g., having different flavors) and, therefore, having a different reward effect. Theorem A3 predicts that, if the session duration is long enough, early in the session the response rate for the outcome with the greater desirability will be higher whereas, later in the session, responses for the other outcome will increase.

## Discussion

Computational models of action selection are essential for understanding decision making processes in humans and animals, and here we extended them by providing a general analytical solution to the problem of response vigor and choice. There are two significant differences between the model proposed here and previous models of response vigor (Dayan, 2012; Niv et al., 2007). Firstly, although the effect of between-session changes in outcome values on response vigor was addressed in previous models (Niv, Joel, &Dayan, 2006),the effects of on-line changes in outcome values within a session were not addressed. On the other hand, the effect of changes in outcome value on the choice between actions has been addressed in some previous models (Keramati & Gutkin, 2014), however their role in determining response vigor has not been investigated. We address such limitations directly in this model.

Secondly, previous work conceptualized the structure of the task as a semi-Markov decision process and derived the optimal actions that maximize the average reward per unit of time (average reward). Here, we used a variational analysis to calculate the optimal actions that maximize the reward earned within the session. One benefit of the approach taken in the previous works is that it extends naturally to a wide range of instrumental conditioning schedules such as interval schedules, whereas the extension of the model proposed here to the case of interval schedules is not trivial. Optimizing the average reward (as adopted in previous work) is equivalent to the maximization of reward in an infinite-horizon time scale; i.e., the session duration is unlimited. In contrast, the model used here explicitly represents the duration of the session which, as we showed, plays an important role in the pattern of responses.

**
*Relationship to principle of least action.***
 A basic assumption that we made here is that the decision-maker takes actions that yield the highest amount of reward. This reward maximization assumption has a parallel in physics literature known as *the principle of least action*, which implies that among all trajectories that a system can take, the true trajectories are the ones that minimize the action. Here action has a different meaning from that used in psychology literature, and it refers to the integral of the Lagrangian (*L*) along the trajectory. In the case of a charged particle with charge *q* and mass *m* in a magnetic field *B*, the Lagrangian will be:

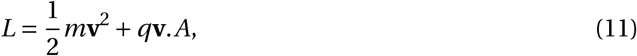
where *A* is the vector potential (*B* =∇ × *A*). By comparing equation 11 with equations 4 and 5, we can see that the reward field *A*_**x**, *t*_ corresponds to the vector potential, and the term *K*_**v**_ corresponds to 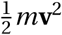 by assuming *m* = 2*ak*
^2^, and *b* = 0. This parallel can provide some insights into the properties of the response rates. For example, it can be shown that when the Lagrangian is not explicitly dependent on time (time-translational invariance), which here implies that 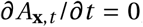, then the Hamiltonian (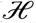, or energy) of the system is conserved. The Hamiltonian in the case of the system defined in equation 4 (with single outcome) is:

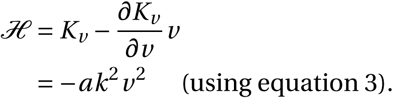

Conservation of the Hamiltonian implies that *ak*
^2^
*v*
^2^ (and therefore *v*) is constant (response-rate is constant), as stated by Theorem 1. Further exploration of this parallel can be an interesting future direction.

## Disclosures and Acknowledgments

A.D completed this work when was employed by CSIRO. B.W.B. was supported by a Senior Principal Research Fellowship from the National Health & Medical Research Council of Australia, GNT1079561.

All authors contributed in a significant way to the manuscript and that all authors have read and approved the final manuscript.

The author declares that the research was conducted in the absence of any commercial or financial relationships that could be construed as a potential conflict of interest.

We are grateful to Hadi Lookzadeh and Peter Dayan for helpful discussions.

## Appendix Value in non-deterministic schedules

The value of a trajectory in the outcome space is the sum of the net amount of rewards that can be earned during a session. However, the amount of reward earned during a session can be non-deterministic, as for example in the case of VR and RR schedules of reinforcement, actions lead to outcomes probabilistically. Similarly, the cost of earning outcomes will also be nondeterministic, since the number of responses required to earn outcomes is non-deterministic. Let’s denote the cost of earning outcomes under such non-deterministic schedules by *K*_**v**_
^′^. Using this, we define the value function as the sum of the *expected* net amount of rewards that will be earned during a session:

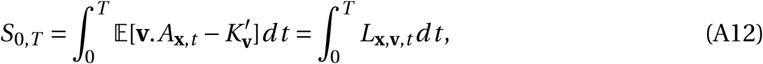
where the expectation is w.r.t the number of earned outcomes along each outcome dimension during *dt* time step. Following the above definition, we have:

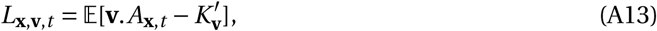
where *L*_**x**,**v**, *t*_ is the expected net reward earned in *dt* time step. In the main text and in the following sections, 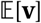 is denoted by **v** for simplicity of notation. By replacing **v** by 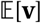 in equation 4 we get:

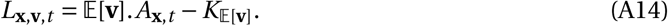

In the main text, equation A14 (equation 4 in the main text) was used instead of equation A13, and the aim of this section is to show that equation A14 and equation A13 are equivalent. We first consider environments with one-dimensional outcome spaces, and then we extend it to the case of environments with multi-dimensional outcome spaces. We maintain the following theorem:

### Theorem A1

Assume that the cost of one response, given that the delay since the last response is τ, is as follows:

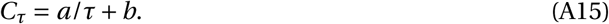

Furthermore, assume that on average, or exactly, k responses are required to earn one unit of the outcome, and r is the number of outcomes earned. Then we have:

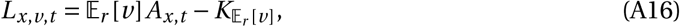

Where

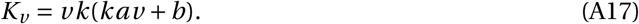

<u>Proof.</u> We first provide an intuitive explanation for why the cost defined in equation A15 is the same as the cost defined in equation A17 in the case of FR schedules of reinforcement (i.e., exactly *k* responses are required to earn an outcome). Earning the outcome at rate *v* implies that the time it takes to earn the outcome is 1/*v*, and since *k* responses have been executed in this period, the delay between responses is:

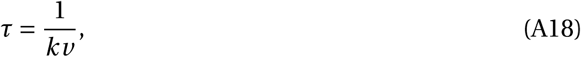
and therefore using equation A15 (equation 2 in the main text), the cost of making one response will be *akv* +*b*. Since *k* responses are required for earning each outcome, the total cost of earning one unit of the outcome will be *k* times the cost of one response, which will be *k*(*akv* +*b*). Since the total amount of outcome earned is *vdt*, the total cost per unit of time will be:

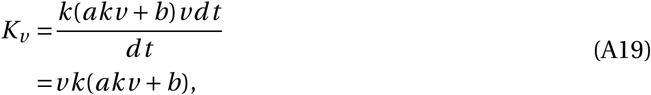

which is equivalent to equation 3 and A17.

We now show that equation A16 and equation A13 are equivalent in order to prove Theorem A1. equation A13 has two terms. As for the first term, *A*_*x*, *t*_ can be assumed to be constant in *dt* time step, and therefore we have:

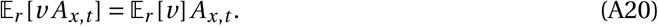

As for the second term we maintain that:

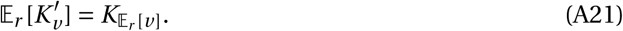

To show the above relation, assume that *r* is the number of outcomes earned after making one Response. Since according to the definition of RR and VR schedules, out of *N* responses on average *N*/*k* will be rewarded, we have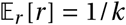 and the expected rate of outcome earning will be:

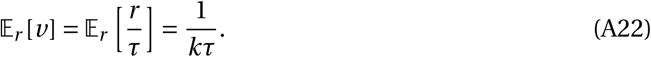

Furthermore, according to equation A15 the cost of one response is *a*/*τ
+b*, and therefore, the cost per unit of time will be:

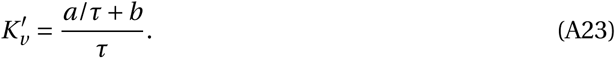

Therefore:

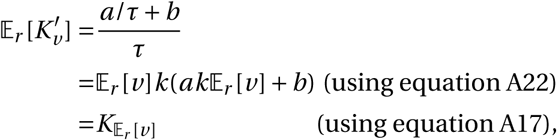

which proves equation A21. Substituting equations A21, A20 in equation A13 yields equation A16, which proves the theorem.

We now turn to the case of multi-dimensional outcome spaces. The aim is to show equation A13 is equivalent to equation A14.To show this, we first maintain that:

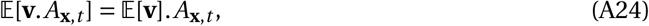

which holds since *A*_**x**, *t*_ can be assumed to be constant during *dt* time step. Next, we show that:

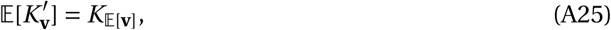

which states that 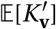 can be represented as a function of 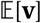. To show this, assume *r*_*i*_ is the number of outcomes earned after making one response for outcome *i*, and *τ*_*i*_ is the delay between responses for outcome *i*. We have:

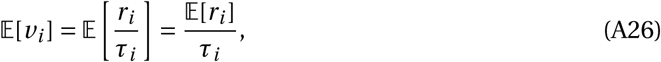

and therefore *τ*_*i*_can be expressed as a function of 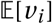. Next, assume that [*C*_*τ*_]_*i*_ is the cost of making one response for outcome *i* with delay *τ*_*i*_ between the responses, and *τ* is a vector containing the delay between responses for each outcome (*τ* = [*τ*_1_… *τ*_*n*_]). In *dt* time step, 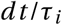 responses for outcome *i* are made, and therefore the total cost in *dt* time period will be [*C*_*τ*_]_*i*_
*dt* /*τ*_*i*_, which implies that the cost for outcome *i* per unit of time is 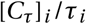. Given this, the total cost paid for all the outcomes per unit of time will be:

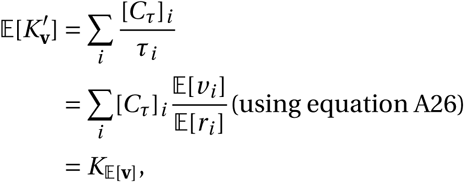

where we used the fact that *τ* in *C*_*τ*_ can be expressed using 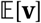 using equation A26, and therefore 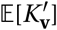 can be expressed as a function of 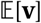, which is denoted by 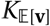(as noted in equation A25). Substituting equation A25, A24 in equation A13 yields equation A14.

**Optimal actions in one-dimensional outcome space**

The aim is to derive optimal actions when the outcome space has one dimension. Given the reward field *A*_*x*, *t*_, the reward of gaining *dx* units of outcome will be *A*_*x*, *t*_
*dx*, and since *dx* = *vdt,* the reward earned in each time step is *v A*_*x*, *t*_. Given that *K*_*v*_ is the cost function (the cost paid in each time step), the net reward in each time step can be written as:

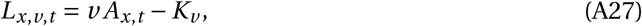

and based on this, the value, which is the sum of net rewards in each time step, will be:

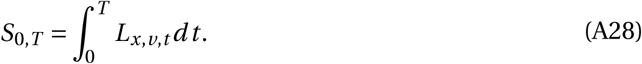

The optimal rates that maximize *S*_0, *T*_ can be found using different variational calculus methods such as the Euler-Lagrange equation, or the Hamilton-Jacobi-Bellman equation (Liberzon, 2011). Here we use the Euler-Lagrange form, which sets a necessary condition for *δS*=0:

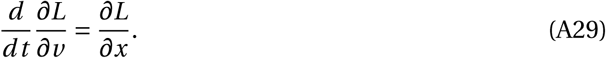

Furthermore, since the end-point of the trajectory is free (the amount of outcomes that can be gained during a session is not limited, but the duration of the session is limited to *T*), the optimal trajectory will also satisfy the transversality conditions:

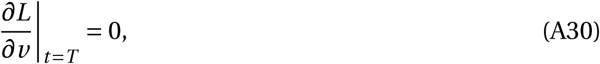

which implies:

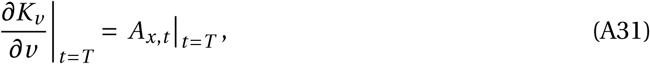

where as mentioned *T* is the total session duration.

By substituting equation A27 in equation A29 we will have:

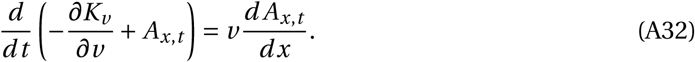

The term 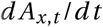 has two components: the first component is the change in *A*_*x*, *t*_ due to the change in *x* and the second component is due to the time-dependent changes in *A*_*x*, *t*_:

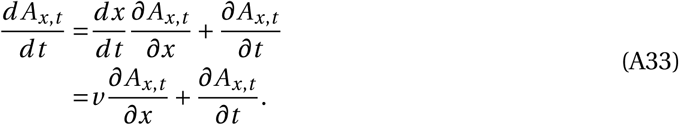

Furthermore we have:

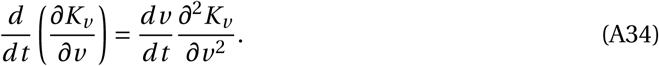

Substituting equations A33, A34 in equation A32 yields:

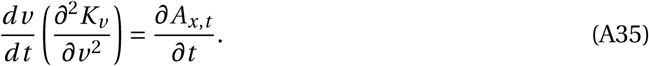

In the condition that the rate of outcome earning is constant (*dv*/*dt* = 0), we have *x*_*T*_ = *vT*, which by substituting in equation A31 yields:

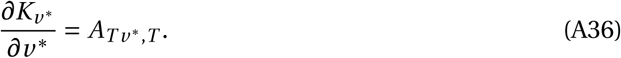

The above equation will be used in order to calculate the optimal rate of outcome earning.

### Theorem 1: Proof

The cost function *K*_*v*_ defined in equation 3 satisfies the following relation:

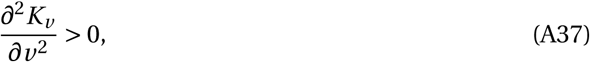

which holds as long as at least one response is required to earn an outcome (*k* > 0), and the cost of earning outcomes is non-zero (*a* > 0).

Assuming that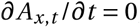, and given equation A37, the only admissible solution to equation A35 is:

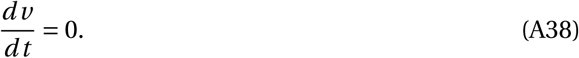

Furthermore, assuming 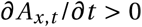, and given equation A37, the only admissible solution to equation A35 is:

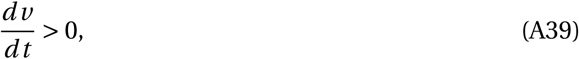
which proves Theorem 1.

### Theorem 2: Proof and simulation details

**Proof of Theorem 2.** In order to prove the theorem, we first provide a lemma. Assuming that the total session duration (*T*) has the probability density function *f* (*T*) and that *f* (*T*) > 0, here we show that the expectation of the total session duration never decreases as time passes throughout the session.

### Lemma 1

*Let’s denote the expectation of the session duration at time t*′ *with T*′

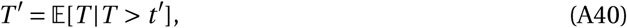
and assume *T* has the following probability density function:

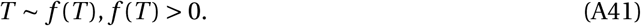

Then:

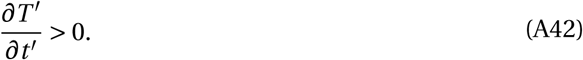

<u>Proof.</u> We have:

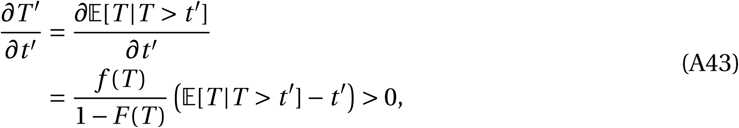
where *F* (*T*) is the cumulative distribution function of *T*.

Based on the above lemma, we show that the optimal response rate is a decreasing function of *t*′ Based on equation A31, the optimal response rate satisfies the following equation:

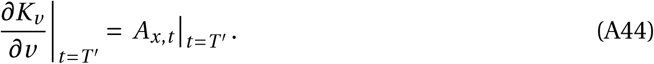

Taking the derivative w.r.t to *t*′ we get:

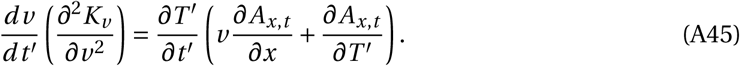

Theorem 2 assumes that 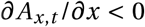 and 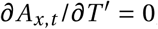, which given equations A42,A37, and

That *v* > 0 yields:

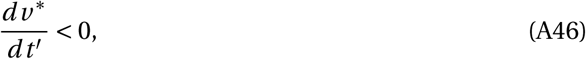

which implies that the rate of earning outcomes decreases as time passes within a session.

**Simulation details.** The simulation of the model depicted in Figure 1: right panel requires defining (i) the reward field, (ii) the cost function, and (iii) a probability distribution over the session duration. As for the probability distribution of the session duration, following McGuire et al (McGuire & Kable, 2013), we assumed that *T* follows a Generalized Pareto distribution:

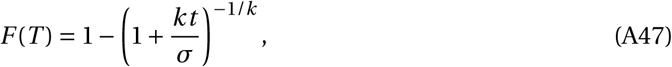

where *k* is a shape parameter (note that *k* is not the ratio-requirement here) and *σ* is a scale parameter, and the third parameter (location *μ*) was assumed to be zero. Furthermore we have:

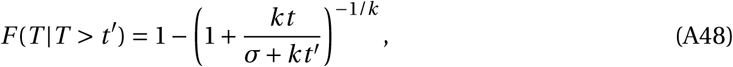

which has the following expected value:

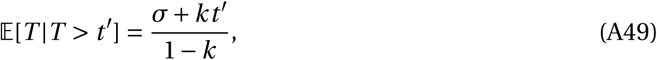

which as we expect is an increasing function of *t′*. For the simulation of the model we assumed that *k* = 0.9 and *σ* = 6, which represents that the initial expectation for the session duration is 60 minutes.

For the cost function, in all the simulations the cost defined in equation 3 was used, which is equivalent to the cost function used in the previous works (Dayan, 2012; Niv et al., 2007).

For the definition of the reward field, we used the framework provided by Keramati et al (Keramati & Gutkin, 2014), which provides a computational model for how the values of outcomes change with the consumptions of the outcomes. They suggested that the dependency of the reward field on the amount of outcome earned is indirect and it is through *the motivational drive*. They conceptualized the motivational drive as the deviations of the *internal states* of a decision-maker from their homeostatic set-points. For example, let’s assume that there is only one internal state, say hunger, where *H* denotes its homeostatic set-point (which corresponds to the deprivation level, assuming that initial value of *x* is zero), and there is an outcome which consuming each unit of it satisfies *l* units of the internal state. In this condition, the motivational drive at point *x*, denoted by *D*_*x*_, will be:

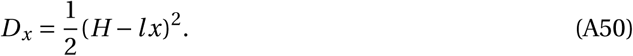

Keramati et al (Keramati & Gutkin, 2014) showed that such a definition of the motivational drive has implications that are consistent with the behavioral evidence. According to the framework, the reward generated by earning *δx* units of the outcome is proportional to the change in the motivational drive, which can be expressed as:

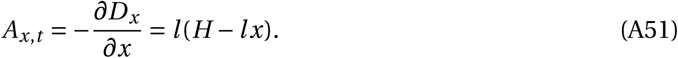

As equation A51 suggests, with earning more outcomes (increase in *x*) *A*_*x*, *t*_ decreases. Given the above reward field, the optimal response rate of outcome earning, obtained by equation A36 will be:

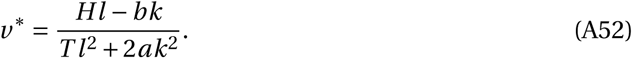

Equation A52 was used in the simulations of the “decreasing reward and unknown session duration” condition in Figure 1: right panel. The simulation of this condition was done using parameters *k* = 15, *l* = 0.1, *a* = 0.002, *b* =0.1, *H* = 80. As for *T*, in each time *t*
^′^ within the session, the expected session duration 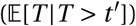 was calculated using equation A49 and was used as *T* in equation A52.

For the “known session duration (fixed or decreasing reward)” condition in Figure 1: right panel, the same parameters as the previous condition were used, but the session duration was fixed to *T* = 60. For the “fixed reward (known or unknown session duration)” condition, we assumed that the reward field is independent of the amount of reward earned:

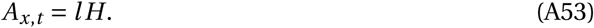

Given the above reward field, the optimal rate of outcome earning is:

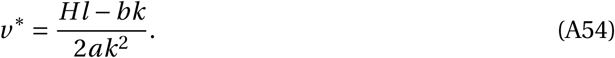

The simulation of this condition was done using parameters *k* = 15, *l* = 0.1, *a* = 0.002, *b* = 0.1, *H* = 40. Note that in this condition the response rate was independent of the session duration. The response rates in all the conditions were obtained by multiplying the outcome rates by *k* (since *k* responses are required to earn one unit of outcome).

**Simulation details of Figures 2 and 3.** The simulation depicted in Figure 2 and Figure 3 are using equation A52 with the following parameters (note that the optimal response rates were obtained by multiplying *v*
^*^ by *k*). For Figure 2: right panel simulation parameters are *T* = 50, *k* = 1, *l* = 1, *a* = 1, *H* = 8. Parameter *b* is varied between 3 to 7 in order to generate the plot.

In Figure 3: left panel simulation parameters are *T* = 50, *l* = 1, *a* = 0.3, *b* = 0.05, *H* = 100.

Parameter *k* was varied between 1 to 100 in order to generate the plot.

In Figure 3: middle panel simulation parameters are *T* = 50, *k* = 1, *l* = 1, *a* = 0.3, *b* = 0.05.

Parameter *H* was varied between 10 to 100 in order to generate the plot.

In Figure 3: right panel simulation parameters are *T* = 50, *k* = 1, *a* = 0.1, *b* = 0.1, *H* = 100.

Parameter *l* was varied between 0 to 1 in order to generate the plot.

**Optimal actions in multi-dimensional outcome space**

The aim of this section is to derive the optimal actions in the condition that the outcome space is multi-dimensional. Optimal trajectory will satisfy the Euler-Lagrange equation along each outcome dimension:

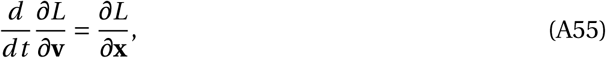

where:

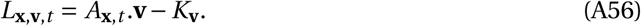

Furthermore since the end point of the trajectory is free (the total amount of outcomes is not fixed) we have:

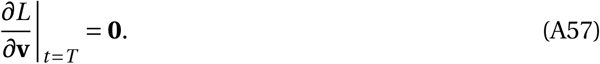

Using equation A55, A56 we have:

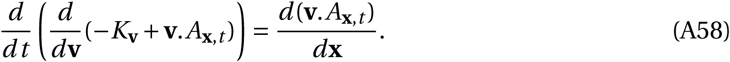

For the right hand side of the above equation we have:

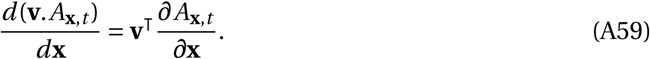

We also have:

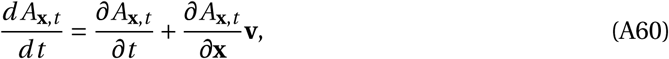

which by substitution into equation A58 yields:

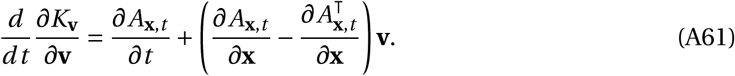

We now provide two lemmas, which will be used in the proof of the following theorems.

### Lemma 2

Assume that **H** is the Hessian matrix of K_**v**_, i.e.,

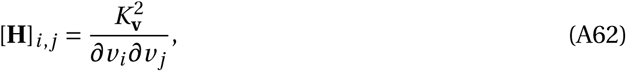
and furthermore assume that the cost of earning outcomes along each dimension is independent of the outcome rate on the other dimensions, i.e.

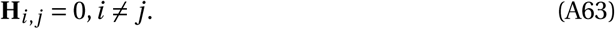

Then:

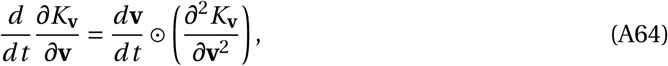
where 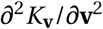 represents the diagonal terms of **H**,and ⊙ is entrywise Hadamard product.

<u>Proof.</u> Using equation A63 we have:

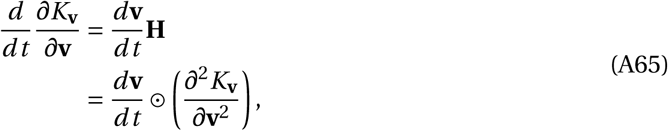
where the last equation comes from the fact that **H** is a diagonal matrix.

### Lemma 3

Assuming that the reward field is conservative, i.e.,

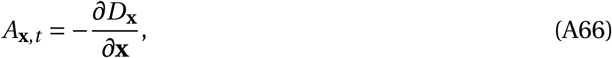
then:

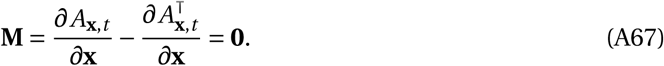

<u>Proof.</u> Using equation A66 we get:

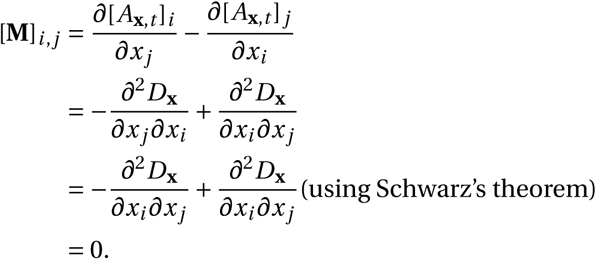

Note that the use of Schwarz’s theorem is based on the assumption that D_**x**_ is twice differentiable, which holds in the circumstances that we consider here.

### Theorem 3: Proof and simulation details

**Proof of Theorem 3.** Theorem 3 assumes that (i) the costs of earning outcomes are independent equation A63,(ii) the reward field is conservative equation A66 and (iii) the reward field is independent of time 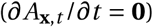. Based on Lemma 2, Lemma 3 and equation A61 we have:

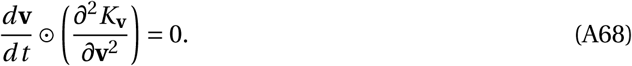

Given that equation A37 holds along each outcome dimension 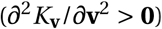, the only admis-sible solution to equation A68 is 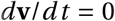, which shows that the optimal rate of earning outcomes is constant. Since the optimal rate is constant, we have **x**_*T*_ = *T*
**v**
^*^, which by substituting in boundary conditions implied by equation A57 yields equation 10:

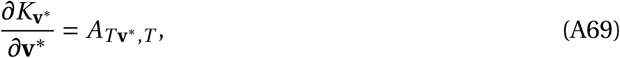
which completes the proof the theorem.

**Simulation details.** For the simulation of the model in Figure 4: left panel “independent cost” condition, it is assumed that the two outcomes have the same reward effect, but earning the second outcome requires *l* times more responses. Following Keramati et al (Keramati & Gutkin, 2014), since the two outcomes have the same reward properties we defined the motivational drive as follows:

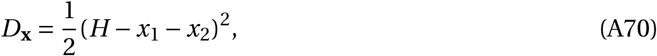
where as mentioned *D*_**x**_ is the motivational drive and it represents the deviations of the internal state of the decision-maker from its homeostatic set-point (*H*). *x*_1_ is the amount of **O**_1_ earned and *x*_2_ is the amount of **O**_2_ earned, and the current motivational drive for earning outcomes depends on the difference between the total amount of earned outcomes (*x*_1_ + *x*_2_) and the homeostatic set-point (*H*).

Given the motivational drive, the amount of reward generated by consuming each outcome will be equal to the amount of change in the motivational drive due to the consumption of the outcomes (equation 8), and therefore, we have:

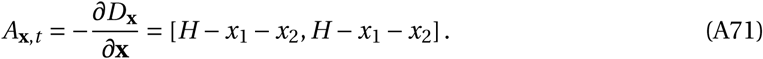

The above equation was used as the reward field in the simulations. As for the cost function, earning one unit of **O**_1_ requires *k* responses on the left lever, and earning one unit of **O**_2_ requires *lk* responses on the right lever. Based on this and using equation 3, the cost function will be:

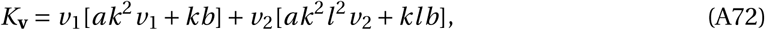
where *v*_1_ is the rate of earning **O**_1_ and *v*_2_ is the rate of earning **O**_2_.

Using Theorem 3, the optimal response rate will be:

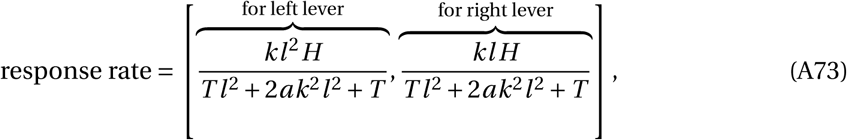
where as mentioned in the main text “left lever” is the response that should be taken for earning **O**_1_, and “right lever” is the response that should be taken for earning **O**_2_. Parameters used for simulations are *k* = 1, *l* = 2, *a* = 1, *b* = 0, *H* = 100.

### Theorem A2: Definition, proof and simulation details

**Proof of Theorem A2.** The aim of this section is to derive optimal actions in the conditions that the costs of earning outcomes are dependent on each other. In this condition, one can assume what determines the cost is the delay between subsequent responses, either for the same or for a different outcome, i.e., the cost is proportional to the rate of earning all of the outcomes. In particular, if for earning **O**_1_, *k* responses are required and for earning **O**_2_, *lk* responses are required (*l* ≠1), then the delay between subsequent responses (*τ*) will be 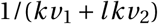. Given equation 2, the cost of earning one unit of **O**_1_ will be *kl*[*a*(*kv*_1_ + *lkv*_2_)+*b*], and the cost of earning one unit of O_2_ will be *kl* [*a*(*kv*_1_ +*lkv*_2_)+*b*]. Such a cost function can be achieved by defining the cost as follows:

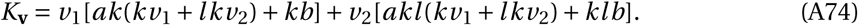

In the following theorem, we maintain that given the above cost function, the optimal actions are to make no response for **O**_2_, and to make responses for **O**_1_ at a constant rate.

### Theorem A2

Given the cost function defined in equation A74 and assuming that the two outcomes have the same reward properties, i.e.:

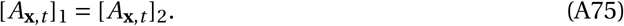

Then the optimal actions satisfy the following equations:

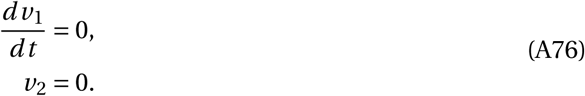

<u>Proof.</u> By substituting equation A74 equation A56 we have:

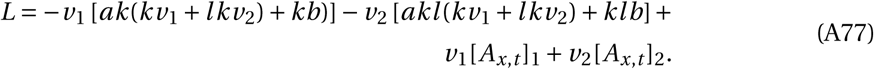

Using the boundary condition mentioned in equation A57 we have:

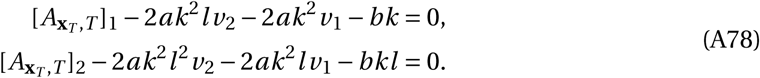

Equation A75 we get:

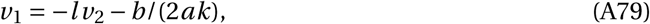
which is not admissible given constraints *v*_1_ ≥ 0 and *v*_2_ ≥ 0, and therefore we assume either *v*_1_ or *v*_2_ will be equal to zero. The trajectory will have a higher value by setting *v*_2_ to zero since **O**_2_ has a higher cost, and therefore the optimal solution will be *v*_2_ = 0. Since *v*_2_ = 0 the problem degenerates to a one-dimensional problem, in which according to Theorem 1 the optimal response rate is constant, and therefore the rate of responding for **O**_1_ will be constant, which proves the theorem.

**Simulation details.** For the simulation of the model in Figure 4: left panel “dependent cost” condition, it is assumed that *k* responses on the left lever are required to earn **O**_1_ and *lk* response are required on the right lever to earn **O**_2_. Similar to the “independent cost” condition mentioned in the previous section, the reward field was assumed as follows:

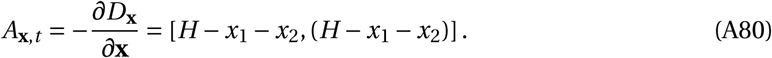

Since the response rate for one of the outcomes will be zero (according to Theorem A2), the problem degenerates to an environment with one action and one outcome. Using Theorem 1, and equation A36 the optimal response rate will be:

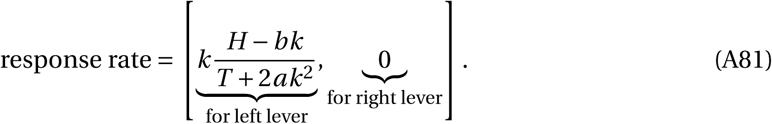

Parameters used for simulations are *k* = 1, *l* = 2, *a* = 1, *b* = 0, *H* = 100.

### Theorem A3: Definition, proof and simulation details

**Proof of Theorem A3.** The aim of Theorem A3 is to derive optimal actions when the reward field is non-conservative and the costs of actions are independent. An example of a nonconservative reward field is when the amount of reward that consuming an outcome produces is greater or smaller than the change in the motivational drive. For example, assume that there are two outcomes available, and the consumption of both outcomes has a similar effect on the motivational drive:

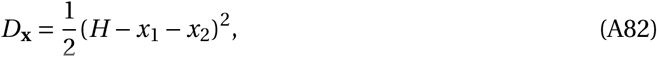
but the reward that the second outcome generates is *l* times larger (*l* ≠1) than the change it creates in the motivational drive:

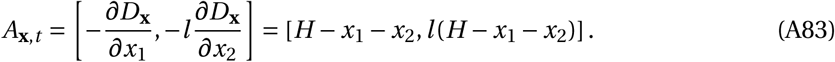

In this condition 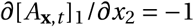 and 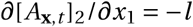, and therefore the reward of the second outcome due to the consumption of the first outcome decreases more sharply than the reward of the first outcome would, due to the consumption of the second outcome. We have:

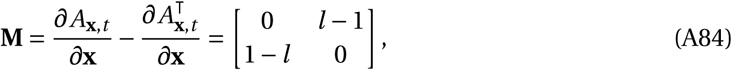
and as long as *l* ≠1 then **M** ≠ **0**, and therefore the reward field is non-conservative, because if it was conservative then according to Lemma 3 we should have **M** = **0**.

If the reward field is non-conservative, i.e., there does not exist a scalar field *D*_**x**_ such that A_**x**, *t*_satisfies equation 8, then the optimal response rates are as follows: early in the session the decision-maker exclusively works for the outcome with the higher reward value (O_1_) and, when the time remaining in the session is less than the threshold (*T*_*c*_), the decision-maker then gradually starts working for the outcome with the lower reward value (O_2_). More precisely we maintain the following theorem:

### Theorem A3

If the reward field follows equation A83, ∂A _x, t_/∂t = **0**, and the cost is as follows:

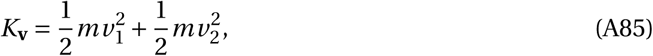
then the optimal trajectory in the outcome space will be:

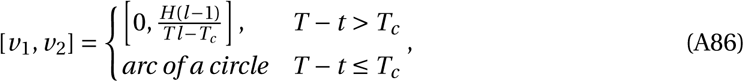
where

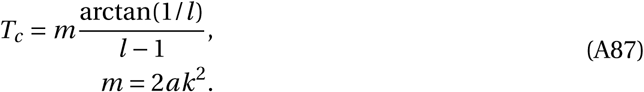

<u>Proof.</u> We have:

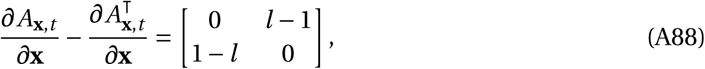
and based on equations A63, A85, A61 we get:

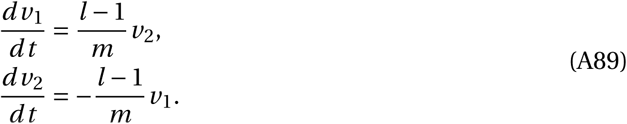

Defining 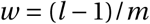, the solution to the above set of differential equations has the form:

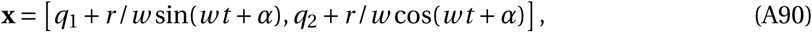
which is an arc of a circle centered at [q_1_, q_2_], and r and α are free parameters. The parameters can be determined using the boundary condition imposed by equation A57, and also assuming that the initial position is **x** = 0. The boundary condition in equation A57 implies:

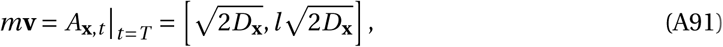
which implies that at the end of the trajectory the rate of earning the second outcome is l times larger than the first outcome. Therefore, the general form of the trajectory will be an arc starting from the origin and ending along the above direction. Given the constraint that **v** ≥ 0 only the solutions in which q_2_ ≤ 0 are acceptable ones (i.e., the center of the circle is below the x-axis). Solving forq_2_ ≤ 0 we get:

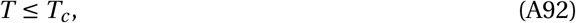
where

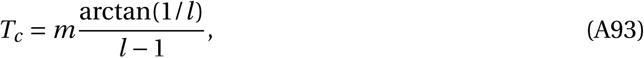
and therefore T_c_ is independent of H (the initial motivational drive). As such if T ≤ T_c_ then the optimal trajectory will be an arc of a circle starting from the origin. Otherwise, if T > T_c_, the optimal trajectory will be composed of two segments. In the first segment, v_1_ will take the boundary condition v_1_ = 0 and the decision-maker earns only the second outcome (the outcome with the higher reward effect). The first segment continues until the remaining time in the session satisfies(the remaining time is less than T_c_after which the second segment starts,which is an arc of a circle defined by. The rate of earning the second outcome,v_2_, in the first segment of the trajectory (when v_1_ = 0) can be obtained by calculating the rates at the beginning of the circular segment. The initial rate at the start of the circular segment is as follows:

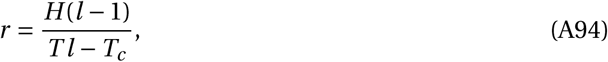
which implies that at the first segment of the trajectory we have:

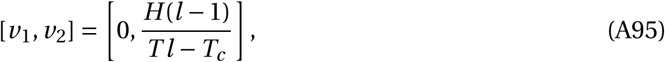
which completes the proof of Theorem A3.

It is interesting to mention that there is a parallel between the trajectory that a decision-maker takes in the outcome space, and the motion of a charged particle in a magnetic field. In the case that the outcome space is three dimensional, using equation A61 the optimal path in the outcome space satisfies the following properties:

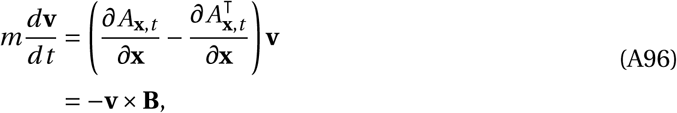
where × is the cross product, **B** is the curl of the reward field (**B** = curlA_**x**, t_), and m = 2ak^2^. The equation A96 in fact lays out the motion of a unit charged particle (negatively charged) with mass m in a magnetic field with magnitude **B**.

**Simulation details.** Simulations shown in Figure 4: right panel are based on Theorem A3, and the parameters used are k = 1, l = 1.1, a = 1, b = 0, H = 100, m = 2ak^2^.

## Footnotes

1 Also known as “First Law of Gossen” named for Hermann Heinrich Gossen (1810 – 1858).

2 Note that in the original notation in (Killeen & Sitomer, 2003), α is denoted by *a* and β is denoted by *b*.

3 Here satiety refers to both post-ingestive factors (such as blood glucose level; (Killeen, 1995)) and/or pre-ingestive factors (for example sensory specific satiety; (McSweeney, 2004))

4 Note that, for simplicity, the simulations in this section are made under the assumption that the session duration is fixed.

